# The nitrogen-fixing fern *Azolla* has a complex microbiome characterized by multiple modes of transmission

**DOI:** 10.1101/2024.05.20.592813

**Authors:** Michael J. Song, Fay-Wei Li, Forrest Freund, Carrie M. Tribble, Erin Toffelmier, Courtney Miller, H. Bradley Shaffer, Carl J. Rothfels

## Abstract

*Azolla* is a floating fern that has closely evolved with a vertically transmitted obligate cyanobacterium endosymbiont—*Anabaena azollae*—that performs nitrogen fixation in specialized *Azolla* leaf pockets. This cyanobacterium has a greatly reduced genome and appears to be in the “advanced” stages of symbiosis, potentially evolving into a nitrogen-fixing organelle. However, there are also other lesser-known inhabitants of the leaf pocket whose role and mode of transmission are unknown. We sequenced 112 *Azolla* specimens collected across the state of California and characterized their metagenomes in order to identify the common bacterial endosymbionts of the leaf pocket and assess their patterns of co-diversification. Four taxa were found across all samples, establishing that there are multiple endosymbionts that consistently inhabit the *Azolla* leaf pocket. We found varying degrees of co-diversification across these taxa as well as varying degrees of isolation by distance and of pseudogenation, which implies that the endosymbiotic community is transmitted by a mix of horizontal and vertical mechanisms, and that some members of the microbiome are more facultative symbionts than others. These results show that the *Azolla* symbiotic community is complex, featuring members at potentially different stages of symbiosis evolution, further supporting the utility of the *Azolla* microcosm as a system for studying the evolution of symbioses.

## Introduction

*Azolla* Lam. (Salviniaceae; Salviniales) is a genus of approximately nine species of floating ferns with specialized leaf pockets that house endosymbiotic bacteria. The most notable microbial inhabitant of the leaf pocket is *Anabaena azollae* (syn. *Nostoc azollae, Trichormus azollae*), which is a nitrogen-fixing cyanobacterium; the *Azolla-Anabaena* symbiosis can fix nitrogen at rates twice that of the legume-*Rhizobium* symbiosis (Watanabe, 1986; Beringer and Johnston, 1984), allowing, among other things, for *Azolla* populations to double their biomass in as little as two days (Peters et al., 1980). The *Anabaena* is vertically transmitted (Zheng et al., 2009) and shows near-perfect codiversification with its *Azolla* host (the *Anabaena* and *Azolla* phylogenies match each other; Li et al., 2018). In addition, the endosymbiont has a considerably reduced genome and can no longer live on its own (Ran et al., 2010; Li et al., 2018): it is potentially evolving into a nitrogen-fixing organelle, like the recently discovered nitroplast in some marine algae (Coale et al., 2024).

There are other known inhabitants of the leaf pocket in addition to the *Anabaena*, including members of the Rhizobiales, two novel species of which have been identified and found to perform denitrification and lack nitrogen fixation genes (Dijkhuizen et al., 2018), and potentially other cyanobacteria like *Fischerella* (Gunawardana and Pushpakumara, 2023). Other studies have further clarified that the *Azolla* microbiome is not only diverse, with many different bacteria having potential roles in promoting plant growth, but also exhibits substantial variation across host species (Banach et al., 2019; Yang et al., 2022). However, there is still no knowledge of the degree to which the members outside of *Anabaena azollae* have co-diversified with the host, and, more generally, the degree to which the symbiotic community is a cohesive entity or an idiosyncratic assemblage of largely independent taxa. A powerful tool for understanding novel associations is examining codiversification (Janz, 2011) between individual endosymbionts and a host. By assessing co-phylogenies and geographic patterns, we can infer whether the members of the core microbiome are mostly obligate or mostly facultative, which will give us a better understanding of the ecosystem within the leaf pocket. In particular, the leaf pocket provides an opportunity to study the evolution of symbioses by allowing glimpses of the process at different stages for different taxa–a spectrum that ranges from loose association and recruitment to intracellular endosymbiosis, such as the chloroplast or nitroplast which represents the far end of this spectrum Coale et al. (2024).

Here, we characterize the metagenome of *Azolla* and assess patterns of co-diversification across the common bacterial endosymbionts, using a large resequencing dataset set generated as part of the California Conservation Genomics Project.

## Methods

### Sample collection and sequencing

#### Field collections and identification

A total of one hundred and twelve samples of *Azolla* were collected (Supplemental Table 1) across the state of California as a part of the California Conservation Genome Project (CCGP; Shaffer et al., 2022). Potential collection sites were identified using the California Consortium of Herbaria Consortium of California Herbaria (2023) and iNaturalist (iNaturalist, 2023). Results were screened for georeferenced observations made after 2016 that included photographs to confirm the *Azolla* identification.

**Table 1:** Testing for isolation by distance. Mantel statistics based on Spearman’s rank correlation rho, permutations = 9999.

| Taxon | Mantel statistic r | Significance |
| --- | --- | --- |
| *Anabaena azollae* | 0.2462 | 1.00E-04 |
| *Rhizobium* | 0.2271 | 0.0017 |
| *Caulobacter* | 0.3791 | 1.00E-04 |
| *Bradyrhizobium* | -0.04252 | 0.7271 |
| *Azolla* Plastome Tree | -0.1013 | 0.9679 |

At each field site, the population was photo-documented at the habitat level and the super-macro level using a Canon PowerShot SX20 IS (Canon, Huntington, New York, USA) and the geographic coordinates were recorded with a Garmin inReach (Garmin Ltd., Schaffhausen, Switzerland). No more than 5% of the total population was sampled, or no more than a 5cm x 5cm mat, whichever was smaller. For plants that were growing neustonically, a pool-skimmer net was used to collect plants away from the shore, in an effort to gather specimens that were relatively undamaged by shoreline wave action. Plants growing terrestrially were carefully removed from the substrate as a matt. All specimens were placed into small, sealable plastic containers and stored in a cooler with ice until they were processed. If collections were made from the same water body, they were taken from at least 1 km apart.

#### Surface sterilization

In order to remove any surface bacterial contaminants, we followed a modified version of the surface sterilization the protocol in Dijkhuizen et al. (2018). After the first treatment, each specimen was transferred to a second 15ml centrifuge tube, and vortexed on low speed three times successively for 1–2 seconds in 3–4ml deionized water. Following the rinse, each individual was surface-sterilized by placing it into a 15ml centrifuge tube with 3–4ml 10% bleach and vortexing on low speed for 3–4 seconds. Once surface sterilized, the specimens were rinsed as above, dried with kimwipes, and flash-frozen at -80°C until it was time to extract DNA.

#### DNA extraction and sequencing

Total genomic and symbiont DNA was extracted using using standard CTAB extraction protocols described in (Doyle and Doyle, 1987). After extractions were complete, DNA quality and quantity were checked using 1% Agarose gel eletrophoresis with 0.1% GelRed Nuclease Dye (Biotum Inc., Fremont CA, USA) and Qubit spectrometry (Thermo Fisher Scientific, Waltham, Massachusetts, U.S.).

Library preparation was performed by the QB3-Berkeley Functional Genomics Laboratory at UC Berkeley. DNA was fragmented with an S220 Focused-Ultrasonicator (Covaris), and libraries prepared using the KAPA Hyper Prep kit for DNA (Roche KK8504). Truncated universal stub adapters were ligated to DNA fragments, which were then extended via PCR using unique dual indexing primers into full length Illumina adapters. Library quality was checked on an AATI (now Agilent) Fragment Analyzer. Libraries were then transferred to the QB3-Berkeley Vincent J. Coates Genomics Sequencing Laboratory, also at UC Berkeley. Library molarity was measured via quantitative PCR with the KAPA Library Quantification Kit (Roche KK4824) on a BioRad CFX Connect thermal cycler. Libraries were then pooled by molarity and sequenced on an Illumina NovaSeq 6000 S4 flow-cell for 2 x 150 cycles, targeting at least 10Gb per sample. Fastq files were generated and demultiplexed using Illumina bcl2fastq2 v2.20 and default settings, on a server running CentOS Linux 7. Whole genome resequencing was performed at an average read depth of around 10x (mean = 12.16x; standard deviation = 5.9; min = 3.6x; max = 44.3x).

Additionally, one reference sample (collection FF365) was sequenced using PacBio Hifi Sequencing (Hifi) producing long-read sequencing data (average depth = 65.56x). The HiFi SMRTbell library was constructed using the SMRTbell gDNA Sample Amplification Kit (Pacific Biosciences, Menlo Park, CA; Cat. no. 101-980-000) and the SMRTbell Express Template Prep Kit 2.0 (Pacific Biosciences; Cat. no. 100-938-900) according to the manufacturer’s instructions. Approximately 10 kb sheared DNA by the Megaruptor 3 system (Diagenode, Belgium; Cat. no. B06010003) was used for removal of single-strand overhangs at 37C for 15 minutes, DNA damage repair at 37C for 30 minutes, end-repair and A-tailing at 20C for 30 minutes and 65C for 30 minutes, and ligation of overhang adapters at 20C for 60 minutes. To prepare for library amplification by PCR, the library was purified with ProNex beads (Promega, Madison, WI; Cat. no. NG2002) for two PCR amplification conditions at 15 cycles each then another ProNex bead purification. Purified amplified DNA from both reactions were pooled in equal mass quantities for another round of enzymatic steps that included DNA repair, end-repair/A-tailing, overhang adapter ligation, and purification with ProNex beads. The PippinHT system (Sage Science, Beverly, MA; Cat no. HPE7510) was used for SMRTbell library size selection to remove fragments *<* 6–10 kb. The 10–11 kb average HiFi SMRTbell library was sequenced at UC Davis DNA Technologies Core (Davis, CA) using one 8M SMRT cell, Sequel II sequencing chemistry 2.0, and 30-hour movies each on a PacBio Sequel IIe sequencer.

All data generated by CCGP can be found in the NCBI SRA (PRJNA720569).

### Metagenome assembly and binning

We mapped the HiFi reads from our reference-genome sample against the *Azolla filiculoides* genome (v1.2; Li et al., 2018) using BWA v0.7.17 (Li and Durbin, 2010); this *A. filiculoides* genome was sequenced from an axenic, symbiont-free strain. We then extracted the unmapped reads using Samtools v1.9 (Li et al., 2009). These unmapped reads, which we expect are from the metagenome, were then uploaded to the BugSeq pipeline (Fan et al., 2021) for long-read taxonomic classification and metagenome binning (Supplemental Table 2). The fasta assemblies from the metagenome binning were used downstream as reference sequences for the leaf pocket microbial taxa. Bacterial genome annotation was performed using Prokka v1.14.6 using default parameters (Seemann, 2014) and genes involved in nitrogen metabolism were identified manually by looking for genes with “nitrogen” in the annotation. The Prokka annotations were used to assess pseudogenation using Pseudofinder v1.1.0 (Syberg-Olsen et al., 2022). Relative abundance of microbes in the resequencing samples was assessed using METAgenomic PHyLogenetic ANalysis for metagenomic taxonomic profiling using default parameters (MetaPhlAn 4.0.3; Blanco-Míguez et al., 2023).

### Variant calling and phylogeny inference

Resequencing samples had Illumina adapters removed and were trimmed using Trimmomatic (v0.39, Bolger et al. (2014)) using the following parameters: LEADING:3 TRAILING:3 MINLEN:36. Samples were then mapped to the *Azolla filiculoides* genome (v1.2; Li et al., 2009) using BWA (0.7.17; Li and Durbin, 2010). bcftools (v1.9; Danecek et al., 2021) was used to create an mpileup for all the samples and to call variants. Variants were filtered to have a QUAL≥30 and to remove multiallelic SNPs and indels, monomorphic SNPs, and SNPs in the close proximity of indels (–SnpGap 10). Reads that did not map to the *Azolla filiculoides* nuclear or chloroplast genome were then recovered and mapped to the metagenome assemblies from the BugSeq output: *Anabaena azollae, Rhizobium, Caulobacter, Bradyrhizobium*, and *Rhizobiaceae*. For each microbial taxon, an mpileup was created and variants were called using bcftools (v1.9, Danecek et al. (2021)) with the following additional parameter: –ploidy 1. Variants were then filtered to have a QUAL≥30 and no monomorphic SNPs using bcftools (v1.9; Danecek et al., 2021).

Alignments of SNPs for the *Azolla* samples and for each of the five focal microbionts that were found in each sample (*Anabaena azollae, Rhizobium, Caulobacter, Bradyrhizobium*, and *Rhizobiaceae* sp. 2) were then created using the phylo command in VCF-kit (v0.2.9; Cook and Andersen, 2017)). For the *Azolla* alignment, heterozygous sites were excluded as per the recommended usage (Cook and Andersen, 2017). Maximum likelihood trees were then inferred for these taxa using IQTree (v1.6.12; Nguyen et al., 2015) under the GTR+ASC model, which accounts for the variable-only ascertainment bias in SNP datasets, and support values were estimated with the ultrafast bootstrap approximation (Minh et al., 2013; Hoang et al., 2018) with 1000 bootstrap replicates.

Assemblies and alignments of whole *Azolla* chloroplasts were produced using GetOrganelle v1.7.7.0 and Homblocks, both with default parameters (Bi et al., 2018; Jin et al., 2020). We included in this alignment all of the previously published *Azolla* chloroplast genomes (Genbank: MF177092.1, MF177091.1, ON684377.1, MF177090.1, MF177089.1, MF177088.1, MF177094.1, MF177093.1). From this alignment we inferred a tree using IQTree v1.6.12 under a GTR+I+G model with 1000 ultrafast bootstrap replicates (Nguyen et al., 2015).

We also used a coalescent-based approach, since we expect incongruence due to our intraspecific sampling (Supplemental Figure 4). Trees were inferred from the *Azolla* nuclear SNP alignment described above using SVDquartets (Chifman and Kubatko, 2014) implemented in PAUP* (v4.0; Swofford, 2003). One hundred bootstrap replicates were performed with quartet sampling of 100,000 quartets. We then made a majority rule consensus tree from the bootstrap trees and resolved polytomies randomly for downstream analysis that require bifurcating trees using the “fix.poly” function in the RRphylo package in R (Castiglione et al., 2018).

### Codiversification and spatial analyses

Codiversification analyses and data visualization was performed using R (R Core Team et al., 2013) with the packages ape (Paradis et al., 2004) and phytools (Revell, 2012). For the plastome tree, we rooted the tree with *Azolla nilotica*. The other trees were visualized using midpoint rooting since they are unrooted; this orientation resulted in two major clades—*A. filiculoides* + *A. rubra* on one side and *A. caroliniana, A. microphylla*, and allies on the other—consistent with the results of studies that applied outgroup rooting (*e*.*g*., Li et al., 2018; Metzgar et al., 2007; Madeira et al., 2013). Statistical tests of codiversification were performed between two trees based on tree distance for both RF (Robinson and Foulds, 1981) and SPR (Swofford, 1990) distances using the “cospeciation” function in phytools. P-values were calculated both by simulation of pure-birth trees and by permutation of tip labels on a fixed tree. Five hundred simulations (and 500 permutations) were performed for each test of the null hypothesis that there are no similarity between trees. Another statistical test of codiversification was performed using the “parafit” function (described in Legendre et al., 2002) in ape using the default parameters and 999 permutations.

The spatial distribution of the taxa was visualized using R and the packages phytools (Revell, 2012) and maps (code by Richard A. Becker et al., 2021). A Mantel test based on Spearman’s rank correlation rho was performed with 9999 permutations using the vegan package in R in order to test for isolation by distance (Oksanen et al., 2022).

## Results

### Characterization of the *Azolla* leaf pocket metagenome

From the reference-genome HiFi data, the BugSeq pipeline assembled 30 unique metagenomic bins ranging from 7.1 Kbp to 5,361.4 Kbp with N50 scores ranging from 7.1 to 1,079.5 Kbp (Supplemental Table 2). Five of these putative bacterial genomes were found in all of our samples. Two of these genomes were nearly complete according to BUSCO scores using the default BugSeq pipeline datasets (Simão et al., 2015; Fan et al., 2021): *Anabaena azollae* and *Rhizobium* sp. Genomes with mapped reads from every sample were included for downstream analysis. These included *Anabaena azollae* (53.6%), *Rhizobium* sp. (1.7%), *Bradyrhizobium* sp. (0.2%), *Caulobacter* sp. (0.2%) and a member of Rhizobiaceae species 2 (0.2%). When comparing the phylogenies inferred from SNP data, we found that the trees of *Rhizobium* sp. and Rhizobiaceae sp. 2 were essentially identical, implying that they are the same taxon and therefore, only *Rhizobium* sp. was analysed in the later analyses.

The prokka annotation found six nitrogen fixation (nif) coding sequences in *Anabaena azollae* and none in *Caulobacter* sp., *Rhizobium* sp., or *Bradyrhizobium* sp. Two *nar* nitrate reductase coding sequences were found in *Anabaena azollae* and none in *Caulobacter* sp., *Rhizobium* sp., or *Bradyrhizobium* sp. No *nir* nitrite reductase or *nos* nitrous oxide reductase coding sequences were found in any of the assemblies.

The Pseudofinder analysis revealed that there is some pseudogenization occuring across all of the common microbiome members. In the main endosymbiont, 32.4% of genes were found to be pseudogenized; *Rhizobium* sp. was found to have 5.6% of genes pseudogenized, *Bradyrhizobium* sp. 6.3%, and *Caulobacter* sp. 5.8%.

The MetaPhlAn taxonomic profiling of samples revealed that the vast majority of reads were of the main endosymbiont cyanobacterium (average across all samples = 94.89%, Supplemental Figure 2) followed by Proteobacteria (4.6%). However, for two samples (FF0505, FF0489) *Anabaena azollae* was not found to be at the highest abundance, possibly due to contamination or disease. A diverse group of phyla were found at much lower abundances, including: Actinobacteria, Armatimonadetes, Bacteroidetes, Chloroflexi, Cyanobacteria, Firmicutes, Gemmatimonadetes, Nitrospirae, Planctomycetes, Proteobacteria, Spirochaetes, Verrucomicrobia, as well as unclassified bacteria.

### *Azolla* phylogenies

Phylogenies inferred from the plastome sequences (Supplemental Fig. 1), the concatenated nuclear SNPs (Supplemental Fig. 3), and the nuclear “species tree” inferred with SVDquartets (Supplemental Figure 4) were all congruent. For simplicity, and to avoid any complications due to coalescent variance, we focus down-stream analyses on the plastome phylogeny. Most distantly related to the other samples is a clade corresponding to *A. filiculoides* (Clade 3, Figure 1). This *A. filiculoides* group has surprisingly very long branch lengths on the concatenated-SNP tree (Supplemental Fig. 3). Our other samples fall in two clades, the largest of which (Clade 2, Figure 1) includes reference sequences of *A. caroliniana* and the smaller of which (Clade 1, Figure 1) is sister to (but divergent from) published plastomes of *A. mexicana/microphylla* (Supplemental Fig. 1; a putative *A. pinnata* sequence that falls in this last clade is an “UNVERIFIED” sequence on Genbank that is almost certainly misidentified—*Azolla pinnata* belongs to a different sub-genus and is only distantly related to the taxa in this study).

**Figure 1:**
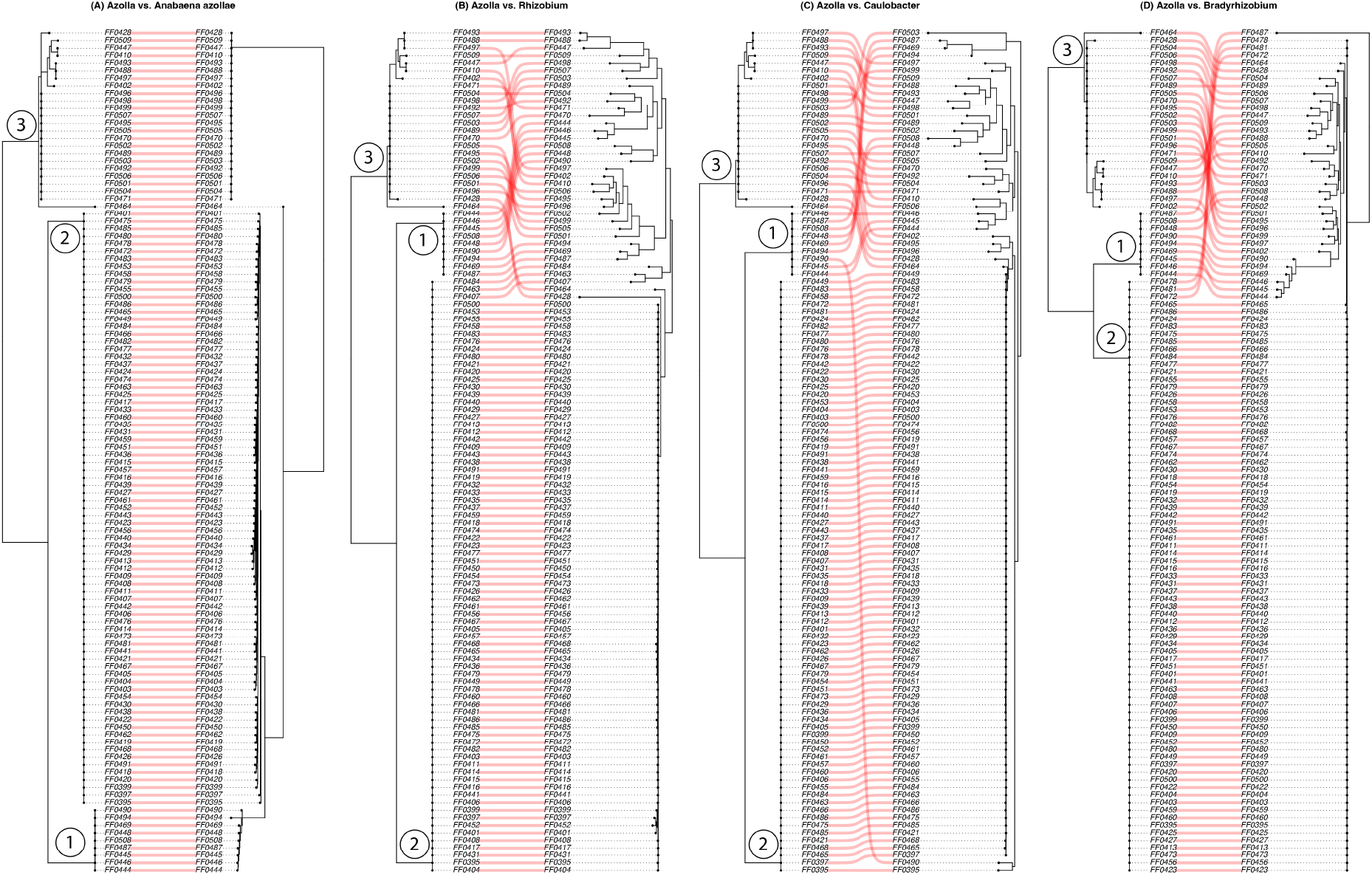
Co-phylogenies in order from left to right: Azolla and *Anabaena azollae, Azolla* and *Rhizobium, Azolla* and *Caulobacter*, and *Azolla* and *Bradyrhizobium*. All *Azolla* phylogenies are from the plastome data.

### Codiversification of endosymbionts

*Azolla* and its symbionts have varying degrees of co-diversification (Figure 1). *Azolla* and its main endosymbiont *Anabaena azollae* exhibit strong co-diversification, as expected, with three major clades found in each. The other focal symbionts are inferred to have a large major clade that is associated with the Clade 2 in the *Azolla* phylogeny, but with more variation and longer branches in the other more distantly related groups.

The *Azolla* and *Anabaena azollae* trees significantly exhibited co-diversification across four of the five tests (Supplemental Table 3). Similarly, *Rhizobium* sp. also exhibited a strong signal of co-diversification with the plastome tree (4/5 tests significant). The other two taxa (*Caulobacter* sp. and *Bradyrhizobium* sp.) exhibited some signal of co-diversification: 3/5 and 2/5 tests significant, respectively.

The endosymbiont phylogenies also had significant associations with other endosymbionts. *Caulobacter* sp. Was found to be significantly associated with *Bradyrhizobium* sp. across all tests and with *Rhizobium* sp. across four of the five tests. *Bradyrhizobium* sp. was found to be significantly associated with with *Rhizobium* sp. in two tests, and *Anabaena azollae* with *Caulobacter* sp. in two tests and with *Bradyrhizobium* sp. in one test.

### Geographic Patterns

Clades 1 and 2 (red in Supplemental Fig. 5) are generally associated with northern samples and Clade 3 (blue in Supplemental Fig. 5) is frequently southern, although these patterns are not consistent (Supplemental Figure 5). Despite these trends, we cannot reject the null hypothesis that there is no relationship between genetic and geographic distances for *Azolla* (Table 1). This is likewise the case for *Bradyrhizobium*. However for *Anabaena azollae, Caulobacter* sp., and *Rhizobium* sp., there is a significant correlation between the geographic and phylogenetic distance matrices for each taxon, which implies isolation by distance for these groups (Table 1).

## Discussion

### *Azolla* diversity in California

The diversity of *Azolla* in California is often treated under two species: *Azolla filiculoides* Lam. and a second taxon treated as either *Azolla microphylla* Kaulf. or *A. mexicana* C.Presl. (Jepson eFlora, 2020; Flora of North America Editorial Committee, 1993). In addition, the potentially invasive *A. pinnata* has been recently reported from the state (Song et al., 2023). Our results highlight the need for further study of the taxonomic diversity of *Azolla* in California, as we find at least three major clades (within subgenus *Azolla*—*A. pinnata*, in subgenus *Rhizosperma*, is outside our taxonomic scope). Of these three clades, one is fairly confidently attributable to *A. filiculoides* (Supplemental Fig. 1). However, the identity of members of the other two clades is unclear. Surprisingly, neither clade appears to correspond to *A. microphylla/A. mexicana*; apparently, despite our expectation that this would be the most common taxon in California, it was absent from our sample. Instead Clade 1, sister to the *A. microphylla/mexicana* reference sequences, includes samples from eastern North America, and likely corresponds to the taxon usually treated as *A. caroliniana*. If these plants are indeed *A. caroliniana*, that would be a remarkable extension to the known range of that species, which is currently understood to be restricted to eastern North America (Flora of North America Editorial Committee, 1993). Clade 2— comprising 77 of our 112 samples—is even more enigmatic: it is sister to Clade 1 + *A. microphylla/mexicana*, and also corresponds to the taxon usually treated as *A. caroliniana*. Further analysis of just the plastid *trnGR* region, places these sequences within a group containing the lineage Madeira et al. (2013) refer to as an undescribed “*A*. species”. However, a thorough examination of the type specimens and other data will be necessary to confidently attach names to these lineages; this work, and a broader taxonomic revision of *Azolla*, is not within the scope of this paper, but our findings should encourage a thorough reassessment of the group in California given the new genomic resources available.

### The *Azolla* symbiotic community

While dominated by its main endosymbiont, we found three other major endosymbiont taxa in the *Azolla* leaf pocket. *Bradyrhizobium* sp. and *Rhizobium* sp. are both members of Rhizobiales, corroborating the results of Dijkhuizen et al. (2018) who found two major Rhizobiales taxa in the *Azolla filiculoides* leaf pocket. The other member, *Caulobacter* sp., is a well-known and common bacterium in freshwater systems, although it can be found in diverse environments (Wilhelm, 2018). These four taxa were found in every one of the 112 samples. Importantly, we found that all of these non *Anabaena azollae* members exhibit pseudogenization of around 5% of their genes; *Anabaena azollae*, on the other hand, had around 30% of its genes pseudogenized. This high degree of pseudogenization is similar to the main endosymbiont in *Azolla filiculoides*, which is also around 30% pseudogenized (Ran et al., 2010). To put this into context, the normal amount of pseudogenization for non-endosymbiotic bacteria is zero, such as was found in other free-living members of Nostocales (Ran et al., 2010).

When we assessed the phylogenies for co-diversification, we found strong evidence that *Anabaena azollae* is co-evolving with *Azolla*, which is what we expected due to its vertical transmittance (Ran et al., 2010; Li et al., 2018) and underscores the efficacy of this approach. When we assessed the other taxa, we found that there is some signal for co-diversification between *Azolla* and *Rhizobium* sp., *Bradyrhizobium* sp. and *Caulobacter* sp. We also found that the phylogenies of the endosymbionts were associated with each other and hypothesized that this was due to shared patterns of isolation by distance. When we tested for this, we found that the ML plastome tree did not exhibit a pattern of isolation by distance. For the bacterial taxa, all but *Bradyrhizobium* exhibited patterns of isolation by distance.

If everything in the leaf pocket were to be vertically transmitted like *Anabaena azollae*, then all of the phylogenies should mirror that of the host. If nothing is being vertically transmitted, we expect to find random patterns between the endosymbiont and the host and patterns that are geographic. We do not find evidence for either of those scenarios, but rather an intermediate between them, which implies that there is some horizontal transmission as well as some vertical transmission occurring. While the limitations of Mantel tests are well known (Bradburd et al., 2013), this pattern is supported by the additional evidence that these taxa were consistently found in every single sample, they all exhibit considerable degrees of pseudogenation, and that they do not exhibit strong co-diversification with the host.

One potential hypothesis for why there is a consistent core microbiome but only loose patterns of codiversification, could be that some bacteria are able to be consistently recruited (maybe by interfacing with the well characterized ammonium and sucrose exchange; Eily et al., 2019), while others end up in the leaf pocket as a general refugium for bacteria (de Vries and de Vries, 2022). Then once in the leaf pocket, microbes may find themselves being transmitted vertically with the host, but in the absence of strong selection for them, they also may find themselves outcompeted and displaced by similar taxa that invade or that the host also recruits from the environment, only to be picked up again later on.

Even though these were partial genome assemblies and so we were unable to assess all of the genes that these endosymbionts have and metabolic roles that they play, the possibility remains that some taxa have evolved to be freeloaders off of the *Anabaena* nitrogen-fixation symbiosis and its associated metabolic products and may even be in conflict with *Anabaena* (Dijkhuizen et al., 2018). The taxonomy and function of these endosymbionts should be investigated in more depth in the future in order to understand what functions and roles these taxa may play in the leaf pocket, if at all. For example, the symbionts may be in metabolic collaboration, competition, or there may be Black Queen processes operating whereby there is a division of labor (Koskella and Bergelson, 2020; Morris et al., 2012). Future work should attempt to get full length assemblies of these bacterial genomes to assess genome reduction as an additional line of evidence for the differences between these putatively obligate and facultative members of the microbiome.

## Author Contributions

M.J.S, F-W.L., F.F., E.T., C.M., B.S., and C.J.R designed and executed the experiment. M.J.S performed the analyses. M.J.S, F-W.L., F.F., E.T., C.M., B.S., C.M.T. and C.J.R contributed with the manuscript.

## Acknowledgements

We would like to thank Britt Koskella for her discussion and insights.

## Data accessibility

All data generated by CCGP can be found in the NCBI SRA (PRJNA720569).

## Supplemental Tables and Figures

### List of Supplemental Tables

Supplemental Table 1. Sample collection data for this study.

### List of Supplemental Figures

Supplemental Figure 2. Violin plots of the relative abundance of the two most abundant phyla across all samples with the mean and standard deviation in red.

Supplemental Figure 3. ML phylogeny of *Azolla* samples inferred from SNP data with bootstrap values at nodes and branch length units in nucleotide substitutions per site.

Supplemental Figure 4. Coalescent tree inferred using SVDQuartets with bootstrap values at nodes and branch lengths in coalescent units.

---

**Supp. Fig. 1:**
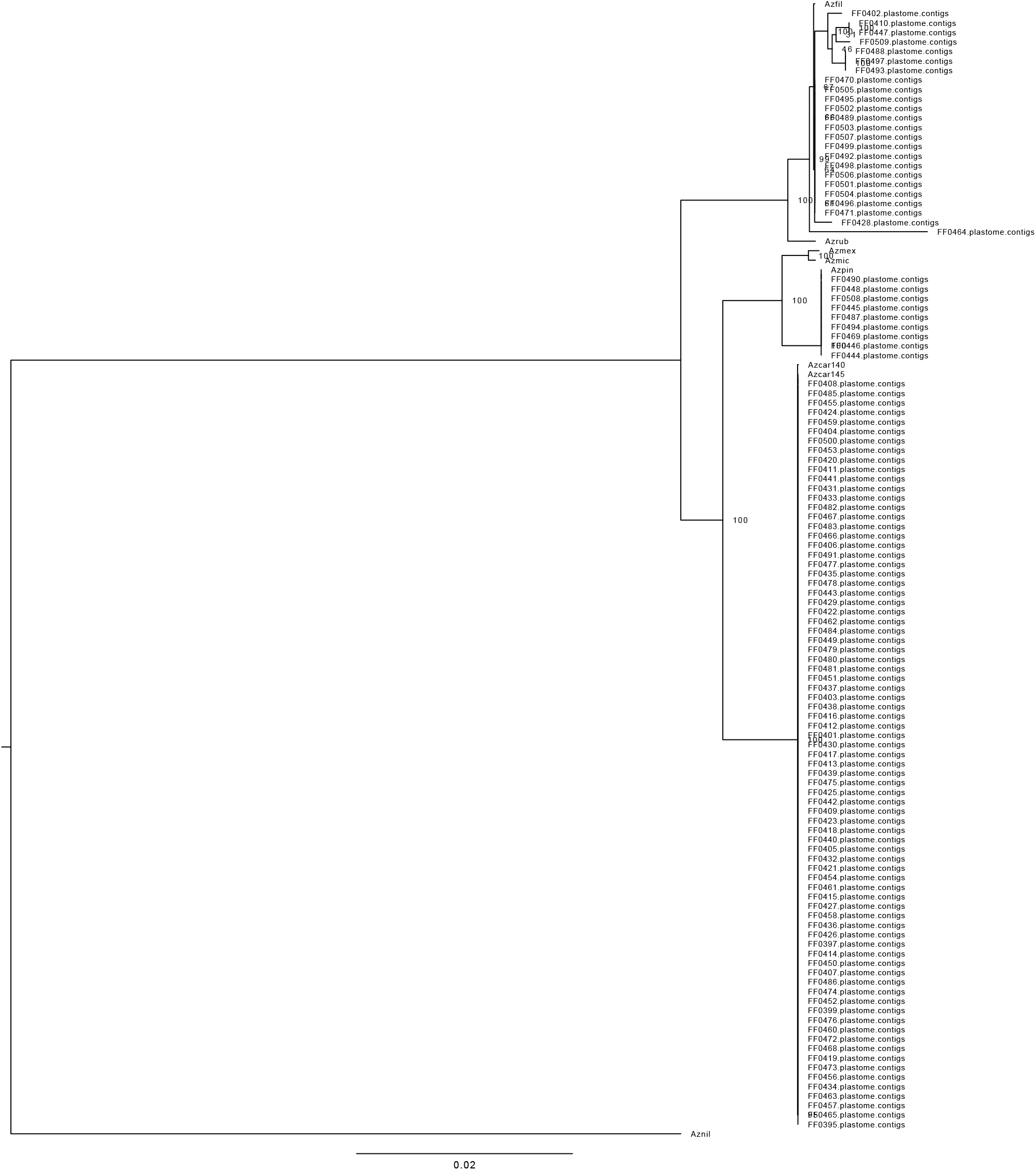
Plastome phylogeny of *Azolla* samples with bootstrap values at nodes. The large clade is found to be similar to *A. caroliniana*, which is sister to a *A. microphyla/A. mexicana* clade (the *A. pinnata* chloroplast being a clear error in Genbank). These two clades are sister to an *A. filiculoides/ A. rubra* clade. These relationships reflect the current understanding of these taxa’s evolutionary history.

**Supp. Fig. 2:**
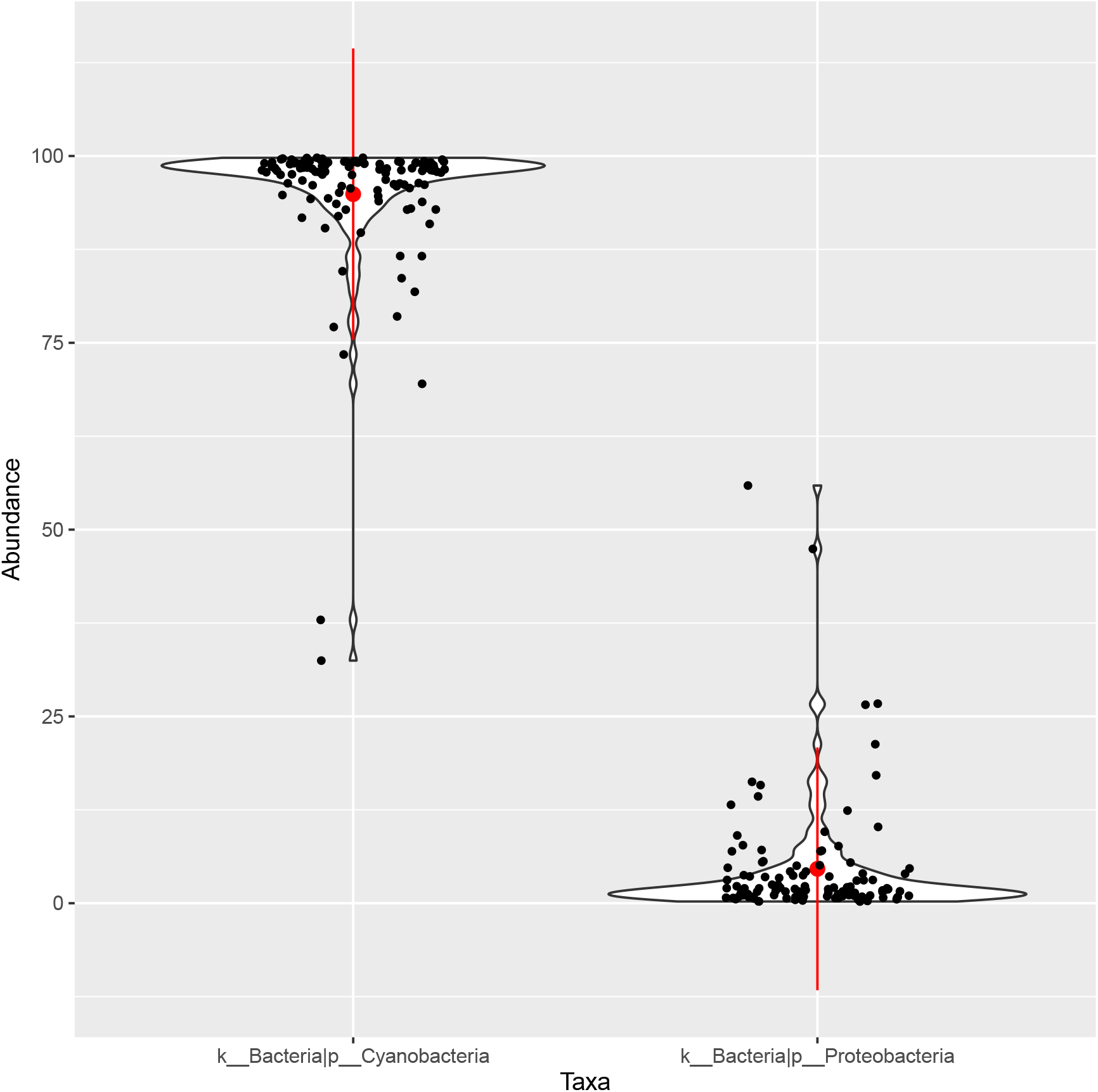
Violin plots of the relative abundance of the two most abundant phyla across all samples with the mean and standard deviation in red.

**Supp. Fig. 3:**
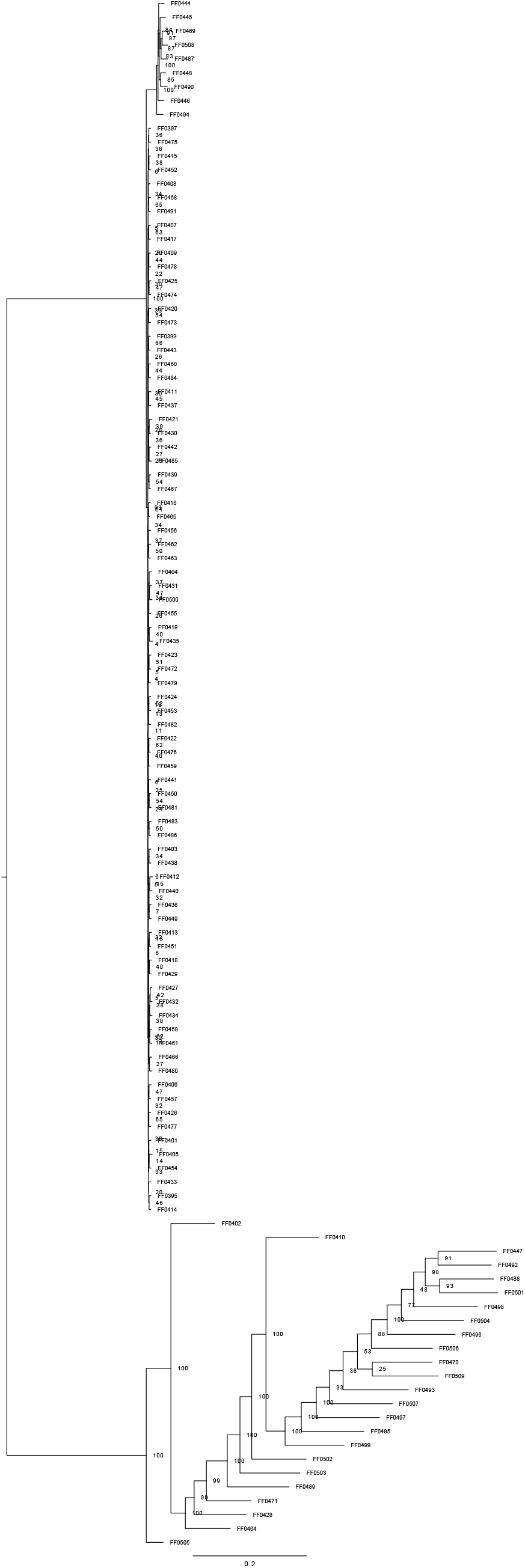
ML phylogeny of *Azolla* samples inferred from SNP data with bootstrap values at nodes and branch length units in nucleotide substitutions per site.

**Supp. Fig. 4:**
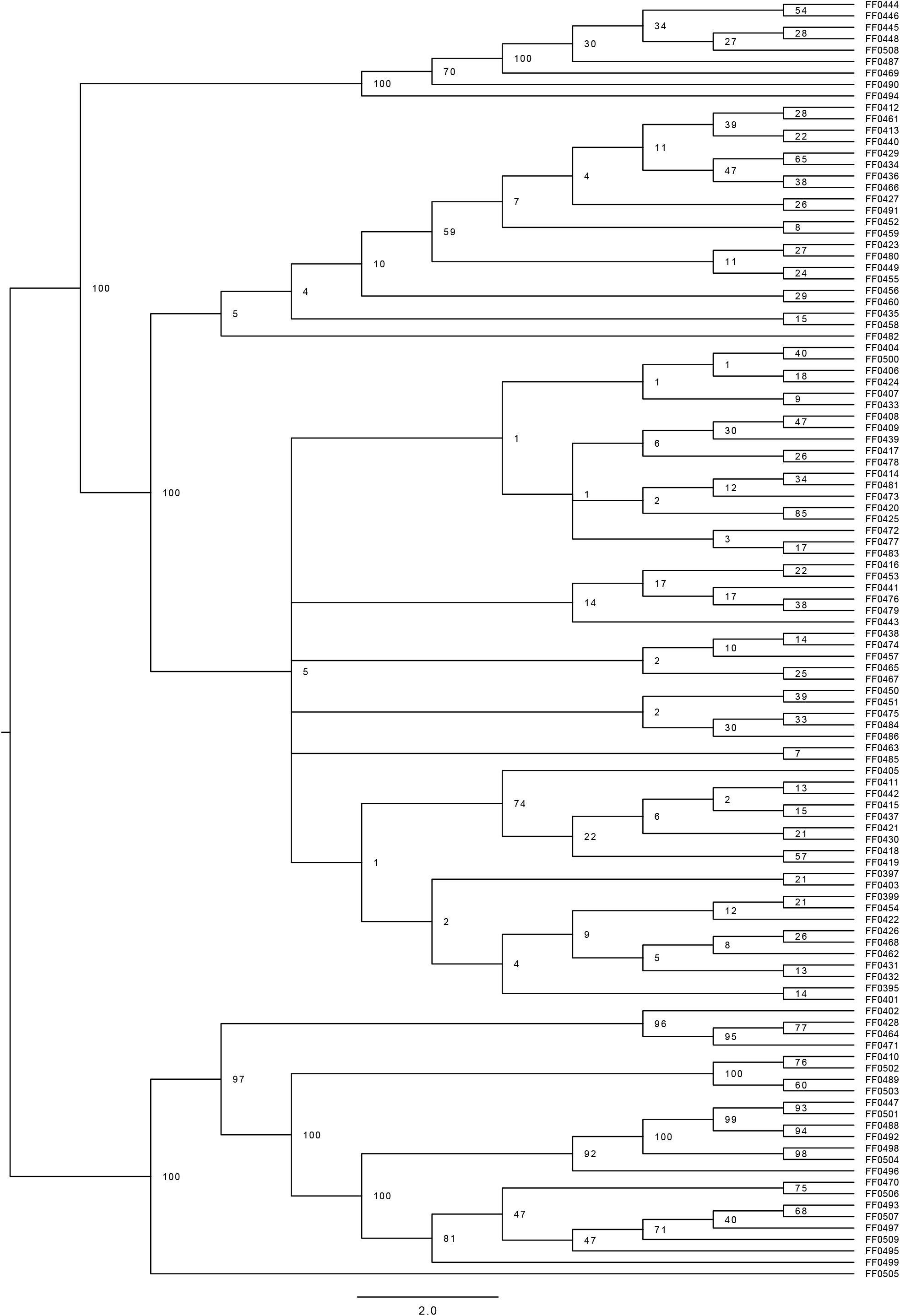
Coalescent tree inferred using SVDQuartets with bootstrap values at nodes and branch lengths in coalescent units.

**Supp. Fig. 5:**
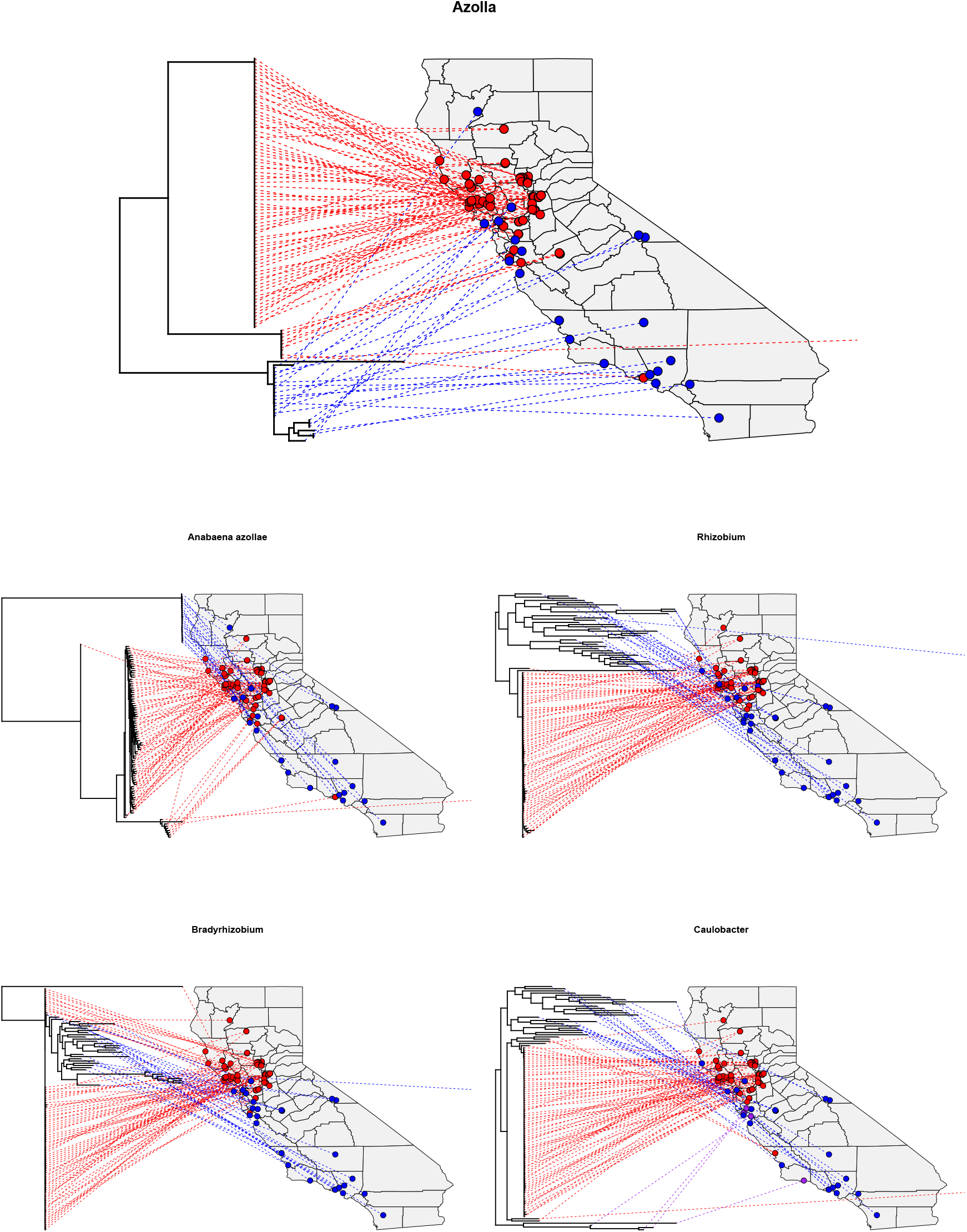
Phylogenies of *Azolla* and its leaf pocket microbial taxa projected onto the samples’ geographic location. Colors are broadly associated across trees to reflect potentially similar major groups in each taxa as well as to assist readability, although they should not be read as formal statements of co-diversification.

**Supp. Tab. 1:**
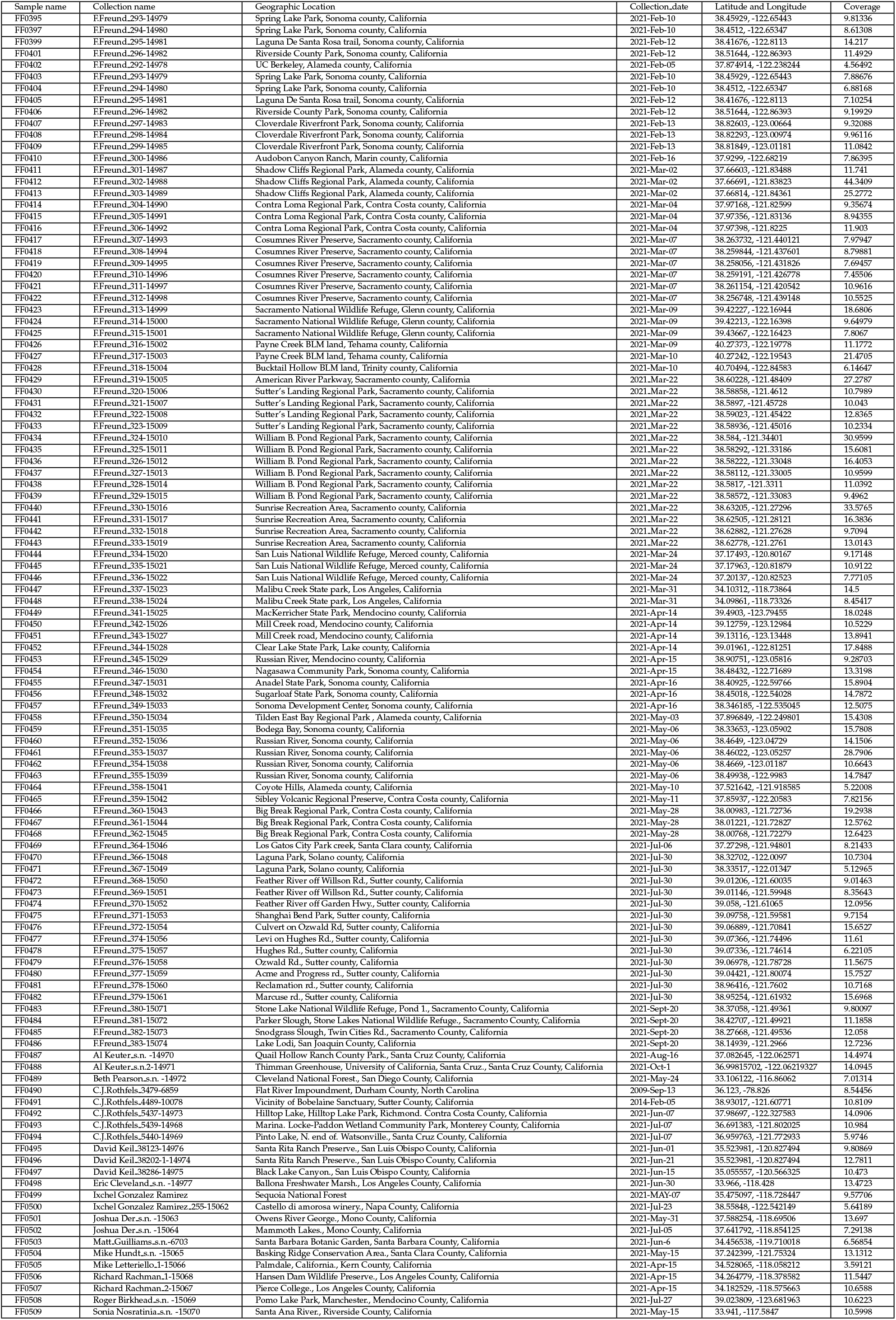
Sample collection data.

**Supp. Tab. 2:** Summary statistics for BugSeq metagenome assembly and taxonomic profiling.

| Sample Name | Abundance | N50 (Kbp) | Assembly Length (Kbp) | ≥ 20X | ≥ 50X | Median | Complete single copy BUSCOs |
| --- | --- | --- | --- | --- | --- | --- | --- |
| *Anabaena azollae* | 53.6% | 1079.5 | 5361.4 | 97.0% | 97.0% | 229.0X | 119 (96%) |
| Root | 32.9% | 26.0 | 39736.7 |  |  |  | NA |
| *Rhizobium* | 1.7% | 451.8 | 4245.0 |  |  |  | 112 (90.3%) |
| *Bradyrhizobium* | 0.2% | 21.5 | 1384.2 |  |  |  | 18 (14.5%) |
| *Caulobacter* | 0.2% | 22.9 | 960.9 |  |  |  | 42 (33.9%) |
| Rhizobiaceae | 0.2% | 304.0 | 516.0 |  |  |  | 9 (7.3%) |
| Alphaproteobacteria | 0.0% | 29.7 | 193.1 |  |  |  | 0 |
| Ascomycota | 0.0% | 11.4 | 11.4 |  |  |  | NA |
| *Bosea* | 0.0% | 8.1 | 8.1 |  |  |  | 0 |
| *Chachezhania* | 0.0% | 10.0 | 10.0 |  |  |  | 0 |
| *Costertonia* | 0.0% | 12.1 | 12.1 |  |  |  | 0 |
| *Dyadobacter* | 0.0% | 7.1 | 7.1 |  |  |  | 0 |
| *Fusarium* | 0.0% | 9.1 | 9.1 |  |  |  | NA |
| *Ginsengibacter* | 0.0% | 11.2 | 11.2 |  |  |  | 0 |
| *Hoeflea* | 0.0% | 12.2 | 12.2 |  |  |  | 1 (0.8%) |
| *Kinneretia* | 0.0% | 7.6 | 7.6 |  |  |  | 0 |
| *Maribacter* | 0.0% | 8.5 | 8.5 |  |  |  | 0 |
| *Methylibium* | 0.0% | 7.4 | 7.4 |  |  |  | 1 (0.8%) |
| *Mucilaginibacter* | 0.0% | 10.2 | 73.7 |  |  |  | 0 |
| *Pandoraea soli* | 0.0% | 8.1 | 8.1 |  |  | 0.0X | 0 |
| *Pedobacter* | 0.0% | 8.7 | 70.5 |  |  |  | 2 (1.6%) |
| *Pirellula* | 0.0% | 8.4 | 8.4 |  |  |  | 0 |
| *Pseudomonadota* | 0.0% | 43.4 | 52.9 |  |  |  | 0 |
| *Pseudoxanthomonas* | 0.0% | 9.1 | 9.1 |  |  |  | 0 |
| *Reyranella* | 0.0% | 9.1 | 9.1 |  |  |  | 1 (0.8%) |
| *Rhodopseudomonas* | 0.0% | 8.8 | 8.8 |  |  |  | 0 |
| Rhodospirillales | 0.0% | 13.9 | 51.6 |  |  |  | 1 (0.8%) |
| *Tardiphaga* | 0.0% | 14.5 | 14.5 |  |  |  | 0 |
| *Trichoderma* | 0.0% | 9.6 | 18.9 |  |  |  | NA |
| *Undibacter* | 0.0% | 7.2 | 7.2 |  |  |  | 0 |

**Supp. Tab. 3:** Statistical tests of codiversification (“cospeciation” in phytools and “parafit” in ape) between two trees based on tree distance. The tests run in the package “cospeciation” were performed for both RF and SPR distances and p-values calculated either by simulation of pure-birth trees (simulation) or permutation of tip labels on a fixed tree (permutation). The number of simulations or permutations for each test: n=500. For the parafit global test, 999 permutations were performed for each test. The null hypothesis for all tests is that there are no similarity between trees.

| Tree 1 | Tree 2 | RF distance permutation, Mean (SD) from null, p-value | RF distance simulation, Mean (SD) from null, p-value | SPR distance permutation, Mean (SD) from null, p-value | SPR distance simulation, Mean (SD) from null, p-value | ParaFitGlobal test statistic, p-value | Tests significant |
| --- | --- | --- | --- | --- | --- | --- | --- |
| *Azolla nuc. tree* | *Anabaena azollae* | 206, 217.7 (0.8), 0.001996 | 206, 216.2 (1.8), 0.001996 | 46, 51.2 (1.1), 0.001996 | 46, 51.1 (1.2), 0.003992 | 24.05071, 0.01 | 5/5 |
| *Azolla nuc. tree* | *Rhizobium* | 218, 217.7 (0.8), 1 | 218, 216.5 (1.6), 1 | 50, 51.1 (1.3), 0.326733 | 50, 51.6 (1.1), 0.138614 | 7.991151, 0.555 | 0/5 |
| *Azolla nuc. tree* | *Caulobacter* | 216, 217.6 (0.8), 0.165669 | 216, 216.4 (1.6), 0.582834 | 40, 50.7 (1.3), 0.0499 | 48, 51.3 (1.1), 0.005988 | 4.79612, 0.735 | 2/5 |
| *Azolla nuc. tree* | *Bradyrhizobium* | 214, 217.6 (0.9), 0.021956 | 214, 214.9 (2.4), 0.457086 | 49, 51 (1.2), 0.103792 | 49, 50.9 (1.2), 0.0998 | 26.48987, 0.001 | 2/5 |
| *Anabaena azollae* | *Caulobacter* | 214, 217.7 (0.8), 0.011976 | 214, 215.1 (2.3), 0.437126 | 49, 50 (1.4), 0.325349 | 49, 50.6 (1.2), 0.181637 | 17.21153, 0.001 | 2/5 |
| *Anabaena azollae* | *Bradyrhizobium* | 212, 217.7 (0.8), 0.001996 | 212, 214.8 (2.4), 0.223553 | 48, 50.3 (1.3), 0.091816 | 48, 50.2 (1.3), 0.093812 | 32.66356, 0.133 | 1/5 |
| *Anabaena azollae* | *Rhizobium* | 216, 217.7 (0.7), 0.127745 | 216, 215.5 (2.1), 0.742515 | 50, 50.4 (1.4), 0.51497 | 50, 50.6 (1.2), 0.461078 | 28.72107, 0.001 | 1/5 |
| *Bradyrhizobium* | *Caulobacter* | 196, 217.7 (0.7), 0.001996 | 196, 213.3 (2.9), 0.001996 | 44, 50 (1.4), 0.001996 | 43, 49.6 (1.4), 0.001996 | 36.28696, 0.312 | 4/5 |
| *Bradyrhizobium* | *Rhizobium* | 204, 217.8 (0.6), 0.001996 | 204, 213 (3), 0.011976 | 48, 50.3 (1.4), 0.111776 | 48, 49.5 (1.3), 0.193613 | 55.31025, 0.233 | 2/5 |
| *Caulobacter* | *Rhizobium* | 217, 217.7 (0.7), 0.001996 | 192, 211.7 (3.4), 0.001996 | 43, 50.1 (1.4), 0.001996 | 43, 49 (1.3), 0.001996 | 18.60991, 0.002 | 5/5 |
| *Azolla SVDQ tree* | *Anabaena azollae* | 200, 217.5 (1), 0.001996 | 200, 213.6 (2.9), 0.001996 | 47, 51.6 (1.1), 0.001996 | 47, 49.9 (1.3), 0.02994 | 67868.59, 0.001 | 5/5 |
| *Azolla SVDQ tree* | *Rhizobium* | 216, 217.7 (0.8), 0.141717 | 216, 214.9 (2.2), 0.832335 | 49, 51.6 (1.1), 0.031936 | 49, 50.7 (1.1), 0.137725 | 24971.56, 0.001 | 2/5 |
| *Azolla SVDQ tree* | *Caulobacter* | 218, 217.6 (0.9), 1 | 218, 215.4 (2.2), 1 | 51, 51.1 (1.2), 0.592814 | 51, 50.6 (1.2), 0.776447 | 16085.25, 0.001 | 1/5 |
| *Azolla SVDQ tree* | *Bradyrhizobium* | 216, 217.6 (0.9), 0.201597 | 216, 215.1 (2.2), 0.818363 | 48, 51.5 (1.1), 0.011976 | 48, 50.6 (1.2), 0.0499 | 39594.57, 0.052 | 3/5 |
| *Azolla plastome tree* | *Anabaena azollae* | 206, 218 (0.2), 0.001996 | 206, 216.2 (1.8), 0.001996 | 39, 41.9 (1.9), 0.103792 | 39, 51.2 (1.1), 0.001996 | 0.02433967, 0.005 | 4/5 |
| *Azolla plastome tree* | *Rhizobium* | 216, 217.9 (0.3), 0.027944 | 216, 216 (1.8), 0.668663 | 39, 41.6 (2), 0.135729 | 39, 51.2 (1.1), 0.001996 | 0.01266035, 0.31 | 4/5 |
| *Azolla plastome tree* | *Caulobacter* | 216, 217.9 (0.4), 0.027944 | 215, 215.7 (2), 0.702595 | 37, 41.4 (2), 0.037924 | 37, 50.9 (1.1), 0.001996 | 0.007746964, 0.694 | 3/5 |
| *Azolla plastome tree* | *Bradyrhizobium* | 218, 217.9 (0.4), 1 | 218, 216.2 (1.7), 1 | 38, 41.7 (2), 0.045908 | 38, 51.1 (1.1), 0.001996 | 0.02544839, 0.664 | 2/5 |

## References

Banach, A., Kuźniar, A., Mencfel, R., and Wolińska, A. (2019). The study on the cultivable microbiome of the aquatic fern azolla filiculoides l. as new source of beneficial microorganisms. Applied Sciences, 9(10):2143.

Beringer, J. E. and Johnston, A. W. (1984). The significance of symbiotic nitrogen fixation in plant production. Critical reviews in plant sciences, 1(4):269–286.

Bi, G., Mao, Y., Xing, Q., and Cao, M. (2018). Homblocks: a multiple-alignment construction pipeline for organelle phylogenomics based on locally collinear block searching. Genomics, 110(1):18–22.

Blanco-Míguez, A., Beghini, F., Cumbo, F., McIver, L. J., Thompson, K. N., Zolfo, M., Manghi, P., Dubois, L., Huang, K. D., Thomas, A. M., et al. (2023). Extending and improving metagenomic taxonomic profiling with uncharacterized species using metaphlan 4. Nature Biotechnology, pages 1–12.

Bolger, A. M., Lohse, M., and Usadel, B. (2014). Trimmomatic: a flexible trimmer for illumina sequence data. Bioinformatics, 30(15):2114–2120.

Bradburd, G. S., Ralph, P. L., and Coop, G. M. (2013). Disentangling the effects of geographic and ecological isolation on genetic differentiation. Evolution, 67(11):3258–3273.

Castiglione, S., Tesone, G., Piccolo, M., Melchionna, M., Mondanaro, A., Serio, C., Di Febbraro, M., and Raia, P. (2018). A new method for testing evolutionary rate variation and shifts in phenotypic evolution. Methods in Ecology and Evolution, 9(4):974–983.

Chifman, J. and Kubatko, L. (2014). Quartet inference from snp data under the coalescent model. Bioinformatics, 30(23):3317–3324.

Coale, T. H., Loconte, V., Turk-Kubo, K. A., Vanslembrouck, B., Mak, W. K. E., Cheung, S., Ekman, A., Chen, J.-H., Hagino, K., Takano, Y., et al. (2024). Nitrogen-fixing organelle in a marine alga. Science, 384(6692):217–222.

code by Richard A. Becker, O. S., version by Ray Brownrigg. Enhancements by Thomas P Minka, A. R. W. R., and Deckmyn., A. (2021). maps: Draw Geographical Maps. R package version 3.4.0.

Consortium of California Herbaria (2023). University of California and Jepson Herbaria. https://ucjeps.berkeley.edu/consortium/. Accessed Mar 3rd, 2023.

Cook, D. E. and Andersen, E. C. (2017). Vcf-kit: assorted utilities for the variant call format. Bioinformatics, 33(10):1581–1582.

Danecek, P., Bonfield, J. K., Liddle, J., Marshall, J., Ohan, V., Pollard, M. O., Whitwham, A., Keane, T., McCarthy, S. A., Davies, R. M., et al. (2021). Twelve years of samtools and bcftools. Gigascience, 10(2):giab008.

de Vries, S. and de Vries, J. (2022). Evolutionary genomic insights into cyanobacterial symbioses in plants. Quantitative Plant Biology, 3:e16.

Dijkhuizen, L. W., Brouwer, P., Bolhuis, H., Reichart, G.-J., Koppers, N., Huettel, B., Bolger, A. M., Li, F.-W., Cheng, S., Liu, X., et al. (2018). Is there foul play in the leaf pocket? the metagenome of floating fern azolla reveals endophytes that do not fix n2 but may denitrify. New Phytologist, 217(1):453–466.

Doyle, J. J. and Doyle, J. L. (1987). A rapid dna isolation procedure for small quantities of fresh leaf tissue. Phytochemical bulletin.

Eily, A. N., Pryer, K. M., and Li, F.-W. (2019). A first glimpse at genes important to the azolla–nostoc symbiosis. Symbiosis, 78:149–162.

Fan, J., Huang, S., and Chorlton, S. D. (2021). Bugseq: a highly accurate cloud platform for long-read metagenomic analyses. BMC bioinformatics, 22:1–12.

Flora of North America Editorial Committee, e. (1993). Flora of North America: Volume 2: Pteridophytes and Gymnosperms, volume 2. Oxford University Press.

Gunawardana, D. and Pushpakumara, B. U. (2023). Molecular characterization of fischerella uthpalarensis, the first subsection v cyanobiont from a tropical azolla species containing dual nitrogenases. bioRxiv, pages 2023–03.

Hoang, D. T., Chernomor, O., Von Haeseler, A., Minh, B. Q., and Vinh, L. S. (2018). Ufboot2: improving the ultrafast bootstrap approximation. Molecular biology and evolution, 35(2):518–522.

iNaturalist (2023). https://www.inaturalist.org/. Accessed Mar 3rd, 2023.

Janz, N. (2011). Ehrlich and raven revisited: mechanisms underlying codiversification of plants and enemies. Annual review of ecology, evolution, and systematics, 42:71–89.

Jepson eFlora (2020). Jepson eflora project.

Jin, J.-J., Yu, W.-B., Yang, J.-B., Song, Y., DePamphilis, C. W., Yi, T.-S., and Li, D.-Z. (2020). Getorganelle: a fast and versatile toolkit for accurate de novo assembly of organelle genomes. Genome biology, 21:1–31.

Koskella, B. and Bergelson, J. (2020). The study of host–microbiome (co) evolution across levels of selection. Philosophical Transactions of the Royal Society B, 375(1808):20190604.

Legendre, P., Desdevises, Y., and Bazin, E. (2002). A statistical test for host–parasite coevolution. Systematic biology, 51(2):217–234.

Li, F.-W., Brouwer, P., Carretero-Paulet, L., Cheng, S., De Vries, J., Delaux, P.-M., Eily, A., Koppers, N., Kuo, L.-Y., Li, Z., et al. (2018). Fern genomes elucidate land plant evolution and cyanobacterial symbioses. Nature plants, 4(7):460–472.

Li, H. and Durbin, R. (2010). Fast and accurate long-read alignment with burrows–wheeler transform. Bioinformatics, 26(5):589–595.

Li, H., Handsaker, B., Wysoker, A., Fennell, T., Ruan, J., Homer, N., Marth, G., Abecasis, G., Durbin, R., and Subgroup, . G. P. D. P. (2009). The sequence alignment/map format and samtools. bioinformatics, 25(16):2078–2079.

Madeira, P. T., Center, T. D., Coetzee, J. A., Pemberton, R. W., Purcell, M. F., and Hill, M. P. (2013). Identity and origins of introduced and native Azolla species in Florida. Aquatic botany, 111:9–15.

Metzgar, J. S., Schneider, H., and Pryer, K. M. (2007). Phylogeny and divergence time estimates for the fern genus Azolla (Salviniaceae). International Journal of Plant Sciences, 168(7):1045–1053.

Minh, B. Q., Nguyen, M. A. T., and Von Haeseler, A. (2013). Ultrafast approximation for phylogenetic bootstrap. Molecular biology and evolution, 30(5):1188–1195.

Morris, J. J., Lenski, R. E., and Zinser, E. R. (2012). The black queen hypothesis: evolution of dependencies through adaptive gene loss. MBio, 3(2):e00036–12.

Nguyen, L.-T., Schmidt, H. A., Von Haeseler, A., and Minh, B. Q. (2015). Iq-tree: a fast and effective stochastic algorithm for estimating maximum-likelihood phylogenies. Molecular biology and evolution, 32(1):268–274.

Oksanen, J., Simpson, G. L., Blanchet, F. G., Kindt, R., Legendre, P., Minchin, P. R., O’Hara, R., Solymos, P., Stevens, M. H. H., Szoecs, E., Wagner, H., Barbour, M., Bedward, M., Bolker, B., Borcard, D., Carvalho, G., Chirico, M., De Caceres, M., Durand, S., Evangelista, H. B. A., FitzJohn, R., Friendly, M., Furneaux, B., Hannigan, G., Hill, M. O., Lahti, L., McGlinn, D., Ouellette, M.-H., Ribeiro Cunha, E., Smith, T., Stier, A., Ter Braak, C. J., and Weedon, J. (2022). vegan: Community Ecology Package. R package version 2. 6–4.

Paradis, E., Claude, J., and Strimmer, K. (2004). Ape: analyses of phylogenetics and evolution in r language. Bioinformatics, 20(2):289–290.

Peters, G. A., Toia Jr, R. E., Evans, W. R., Crist, D. K., Mayne, B. C., and Poole, R. E. (1980). Characterization and comparisons of five n2-fixing azolla-anabaena associations, i. optimization of growth conditions for biomass increase and n content in a controlled environment. Plant, Cell & Environment, 3(4):261–269.

R Core Team, R. et al. (2013). R: A language and environment for statistical computing.

Ran, L., Larsson, J., Vigil-Stenman, T., Nylander, J. A., Ininbergs, K., Zheng, W.-W., Lapidus, A., Lowry, S., Haselkorn, R., and Bergman, B. (2010). Genome erosion in a nitrogen-fixing vertically transmitted endosymbiotic multicellular cyanobacterium. PLoS One, 5(7):e11486.

Revell, L. J. (2012). phytools: an r package for phylogenetic comparative biology (and other things). Methods in ecology and evolution, (2):217– 223.

Robinson, D. F. and Foulds, L. R. (1981). Comparison of phylogenetic trees. Mathematical biosciences, 53(1-2):131–147.

Seemann, T. (2014). Prokka: rapid prokaryotic genome annotation. Bioinformatics, 30(14):2068–2069.

Shaffer, H. B., Toffelmier, E., Corbett-Detig, R. B., Escalona, M., Erickson, B., Fiedler, P., Gold, M., Harrigan, R. J., Hodges, S., Luckau, T. K., et al. (2022). Landscape genomics to enable conservation actions: the california conservation genomics project. Journal of Heredity, 113(6):577–588.

Simão, F. A., Waterhouse, R. M., Ioannidis, P., Kriventseva, E. V., and Zdobnov, E. M. (2015). Busco: assessing genome assembly and annotation completeness with single-copy orthologs. Bioinformatics, 31(19):3210–3212.

Song, M. J., Huynh, M., Lahmeyer, S., and Sedaghatpour, M. (2023). First record of the invasive Azolla pinnata subsp. pinnata (Salviniaceae) in California. American Fern Journal, 113(1):56–57.

Swofford, D. L. (1990). Phylogenetic reconstruction. Molecular systematics, pages 411–501.

Swofford, D. L. (2003). Paup^* phylogenetic analysis using parsimony (^* and other methods). version 4. http://paup.csit.fsu.edu/.

Syberg-Olsen, M. J., Garber, A. I., Keeling, P. J., McCutcheon, J. P., and Husnik, F. (2022). Pseudofinder: detection of pseudogenes in prokaryotic genomes. Molecular biology and evolution, 39(7):msac153.

Watanabe, I. (1986). Nitrogen fixation by non-legumes in tropical agriculture with special reference to wetland rice. Plant and Soil, 90:343–357.

Wilhelm, R. C. (2018). Following the terrestrial tracks of caulobacter-redefining the ecology of a reputed aquatic oligotroph. The ISME journal, 12(12):3025–3037.

Yang, Y.-Q., Deng, S.-F., Yang, Y.-Q., and Ying, Z.-Y. (2022). Comparative analysis of the endophytic bacteria inhabiting the phyllosphere of aquatic fern azolla species by high-throughput sequencing. BMC microbiology, 22(1):1–11.

Zheng, W., Bergman, B., Chen, B., Zheng, S., Xiang, G., and Rasmussen, U. (2009). Cellular responses in the cyanobacterial symbiont during its vertical transfer between plant generations in the azolla microphylla symbiosis. New Phytologist, 181(1):53–61.

